# Different integrins mediate haptotaxis of T lymphocytes towards either lower or higher adhesion zones

**DOI:** 10.1101/509240

**Authors:** X Luo, L Aoun, M. Biarnes-Pelicot, Pierre-Olivier Strale, V Studer, M.-P. Valignat, O. Theodoly

**Affiliations:** LAI, Aix Marseille Univ, CNRS, INSERM, Marseille, France; University of Bordeaux, Interdisciplinary Institute for Neuroscience, Bordeaux, France; CNRS UMR 5297, F-33000 Bordeaux, France; ALVEOLE, Paris, France

**Keywords:** haptotaxis, integrins, chemotaxis, cell guidance, mechanotransduction, lymphocytes, LFA-1, VLA-4

## Abstract

Guidance of cells by molecules anchored on a substrate, known as haptotaxis, is arguably crucial in development, immunology and cancer, however the exact cues and mechanisms driving cell orientation *in vivo* are hardly identified. Adhesive haptotaxis has been described in the case of mesenchymatous cells that develop strong pulling forces with their substrates and orient via a tug of war mechanism – a competition between cells’ pulling edges. In the case of amoeboid cells that migrate with minimal interaction with their substrate, existence of adhesive haptotaxis remains unclear. Here, we studied the crawling of human T lymphocytes on substrates with spatially modulated adhesivity, and observed haptotaxis with surface concentrations of integrin ligands found on high endothelial veinules. Overexpression of ICAM-1 and VCAM-1 molecules observed *in vivo* at transmigration portals can therefore promote leukocyte recruitment. Mechanistically, we show that integrin-mediated haptotaxis of lymphocytes differ both from active chemotaxis, because no mechanotransduction was detected, and from the passive tug of war mechanism of mesenchymatous cells, because different integrins support opposite phenotypes. Cells favored more adherent zones with VLA-4 and, counterintuitively, less adherent zones with LFA-1. These results reveal that integrins control differential adhesive haptotaxis behaviors without mechanotransduction, and this smart capability may support unsuspected ways for cells path selection.

## INTRODUCTION

An efficient immune response relies on the rapid recruitment of circulating leukocytes from the blood system to the inflamed or damaged tissues. Leukocytes first roll and arrest on the endothelium - the cellular monolayer covering the inner lumen of blood vessel - then spread and migrate spontaneously with a rapid crawling mode. Leukocytes eventually cross the endothelium - an event called diapedesis - and reach targeted zones via connective tissues. At vessels inner walls, leukocytes crawl distances of several tens or hundreds of micrometers, proposedly in search for optimal diapedesis sites(1–3). Diapedesis seems indeed to occur at specialized area of the endothelium, called “Cuplike” transmigratory structures(4) or “portals”(3). These structures are composed of vertical microvilli-like projections that are enriched in intercellular adhesion molecule-1 (ICAM-1) and vascular cell adhesion molecule-1 (VCAM-1) and surround transmigrating leukocytes(4). ICAM-1 was found particularly important to favor transmigration in nearby portals(3), which suggests that the endothelium may provide directional guidance to leukocytes at extravasation sites. Guided migration based on adhesion has however never been evidenced for leukocytes.

Directed motion is a hallmark of the immune response. Leukocytes are sensitive to mechanical cues like blood flow (mechanotaxis)(5–8), to soluble biochemical cues like bacterial fragments or chemokines (chemotaxis)(1, 9), and to anchored signaling molecules like chemokines CCL21 or IL8 (haptotaxis)(10–12). However, if directed motion by adhesion molecules (adhesive haptotaxis) is well documented for cancer cells, fibroblasts, endothelial cells or neurons(13–22), it has not been evidenced for leukocytes yet. Hence, adhesive haptotaxis remains largely restricted to mesenchymatous cells. However, mesenchymatous migration is characterized by strong cell/substrate adhesion mediated by mature focal adhesions and strong traction forces transmitted to the substrate by contractile actin stress fibers. Adhesive haptotaxis is therefore directly explained by a tug of war in the cell adherence zone, areas with lower grip destabilizing spontaneously under traction in favor to areas with a higher grip(17). In contrast, amoeboid cells migrate at high speed with low adhesion and low traction forces transferred to the substrate(23–25). A tug of war mechanism, with an interplay of strong adhesion and high traction, is therefore not applicable to amoeboid cells, and the existence of adhesive haptotaxis for leukocytes remains an open question. Furthermore, integrins are much more than adhesion molecules, they can also transduce signals and trigger internal cascades when engaged with their ligands and submitted to a force. Like gradients of anchored chemokines, gradients of integrins ligands could also guide leukocytes via intracellular signaling cascades transmitting molecular detection signals into cytoskeleton reorganization. Two modes of leukocytes guiding by integrin adhesion molecules are therefore plausible, either by active intracellular signaling or by passive competition between adherent cell edges.

To shed light on the existence of integrin-mediated haptotaxis and its physiological relevance and underlying mechanisms, we took advantage of recent technological advances in surface patterning methods(26) to perform controlled *in vitro* experiments. Since leukocytes require a minimal adhesion to resist the shear stress of blood, we prepared adherent substrates with modulations of adhesion properties using the optical molecular printing technique LIMA(26). A series of substrates patterned with stripes of ICAM-1 and VCAM-1 proteins, respectively ligands of integrins LFA-1 (α_L_β_2_) and VLA-4 (α_4_β_1_) were generated to study the crawling of primary human T lymphocytes. Our results demonstrate that crawling lymphocytes display robust integrin-mediated haptotaxis that rely neither on active triggering of intracellular signaling nor on passive tug of war adhesive competition.

## RESULTS

### Protein printing yields substrates patterned with modulated amounts of ICAM-1

The set-up of light induced molecular adsorption LIMA technique(26) uses a 375 nm UV light source coupled optically to an inverted microscope with a Digital Micro-mirror Device (DMD) (**Figure 1**-A). Glass coverslips were first treated with aminopropyltriethoxysilane (APTES), then sequentially incubated with Protein A, Bovine serum albumin (BSA) and chimera Fc-ICAM-1 (**Figure 1**-B). UV illumination was then used to degrade the adhesive properties of ICAM-1 in presence of the photo-activator in a UV-dose dependent manner. Grey-leveled illumination allowed us to obtain substrates with gradual adhesion properties. **Figure 1**-C shows an image of ICAM-1 pattern obtained by immunofluorescence microscopy, where the different fluorescence brightness correspond to different surface concentrations of ICAM-1 molecules. Quantitative analysis of fluorescence images allowed us to calibrate the absolute surface concentration of functional ICAM-1 molecules (**Figure 1**-D).

**Figure 1:**
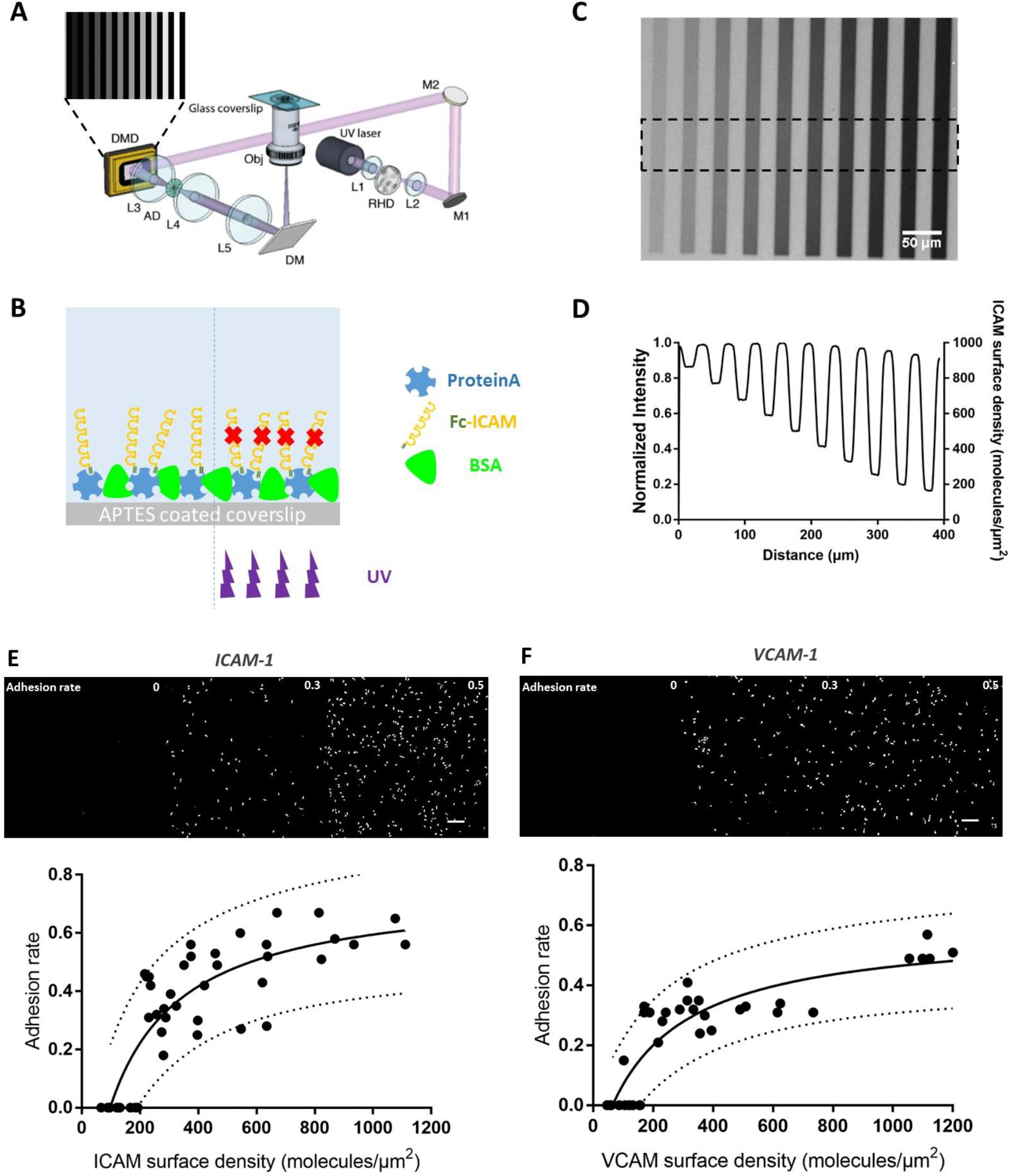
Optical patterning yields substrates with various amounts of grafted integrins ligands and with modulated adhesion. **(A)** Illustration of the optical set-up used for protein patterning (26); **(B)** Illustration of the experimental protocol for ICAM patterning on glass coverslips. Substrates were functionalized with aminosilane APTES, coated with fluorescent protein A, passivated with albumin BSA, then incubated with Fc-ICAM-1 or Fc-VCAM-1, and finally UV-illuminated to modulate the amount of functional integrin ligands. **(C)** Fluorescent images showing patterned stripes with various amounts of protein A. **(D)** The profiles of fluorescence intensity (left axis) and corresponding ICAM-1 surface density (right axis) show decreasing amounts of functional integrin ligands on stripes illuminated with increasing UV-doses. **(E,F) Modulated adhesion versus grafting density**. (Top) Images series showing adhered lymphocytes in presence of a shear flow of 8 dyn/cm^2^ on ICAM-1 (E) and VCAM-1 (F) substrates. (Bottom) Adhesion rate versus ICAM-1 (E) and VCAM-1 (F) surface density. A Langmuir fit was used as a guide for the eye with a confidence interval of 90%.

### Cell adhesion is controlled by surface concentration of integrins ligands

To quantify the effective adhesion of lymphocytes versus the amount of functional ICAM-1 or VCAM-1 molecules anchored on substrates, we prepared homogeneous substrates with various UV-doses allowing different levels of cell adhesion (Figure 1-E and 1-F). Cells were seeded for 10 min on each substrate, then submitted to a gentle shear stress of 1 dyn.cm^−2^ to wash out non-adherent cells. The adhesion strength was then assessed by calculating the ratio between cells still adherent after 5 min under a shear stress of 8 dyn.cm^−2^ and initial adherent cells. This ratio is later called ‘adhesion rate’ (Figure 1-E and 1-F). For ICAM-1 and VCAM-1, the adhesion rate was null below concentrations of 150 -200 molecules.μm^−2^, then increased with concentration until a plateau at 0.6 for concentration around 1000 molecules.μm^−2^ (Figure1-E and 1-F). Resistance to flow is therefore dependent of surface concentration and roughly similar for ICAM-1/LFA-1 or VCAM-1/VLA-4 interactions.

### T cells haptotax on substrates with adherent and non-adherent stripes

A complete characterization of haptotaxis requires a screening of cell guiding phenotype versus two parameters, local concentration and steepness of concentration gradient. The search for the existence of haptotaxis was undertaken here with a single parameter by using step-like profiles made of alternated stripes of different adhesion rates. Cells travelling across adjacent stripes experienced a gradient of infinite steepness, which maximizes conditions for gradient detection by cells. To identify the concentration range where integrins may mediate haptotaxis, we then varied the contrast of adhesion between stripes. The width of stripes was chosen at 20 μm, slightly larger than the average diameter of adherent cells of 15 ± 2 μm, which maximizes the occurrence of simultaneous cell contacts with two adjacent stripes. Alternated stripes of different adhesivity were prepared with a maximum adhesion rate of 0.6 for half of the stripes, and at variable adhesion rate between 0 and 0.6 for the other half. We defined the ‘adhesion contrast’ as the difference in percent between the adhesion rates on alternated stripes normalized over the maximal adhesion rate. An adhesion contrast of 0% corresponds therefore to a homogeneous substrate (with the maximum adhesion rate of 0.6), whereas the adhesion contrast of 100% corresponds to alternated stripes of maximum and null adhesion rates. Cells trajectories were tracked for 15 min and the orientation of each individual cell was assessed from the orientation between first and final positions. Distribution of orientations were displayed on rose wind plots, and to assess directional bias of migrating cells, we further calculated an Anisotropy Index as the difference between the cell fraction in vertical quadrants (orientation bias parallel to stripes) and in horizontal quadrants (orientation bias perpendicular to stripes). In control experiments with homogeneous substrates of maximum adhesion rate 0.6 (**Figure 2**-A and left panel of movie 1 and 2), cells had a random motility, with anisotropy index equal to 0.0 ± 0.1. On alternated stripes of adhesion contrast 100%(**Figure 2**-B and middle panel of movie 1 and 2), cells travelled both on and across adherent and non-adherent stripes. T lymphocytes are indeed able to crawl on adhesive substrates and swim on anti-adhesive substrates(27). However, portions of path with lengths of tens of micrometers appeared with a steady orientation in the direction of stripes (**Figure 2**-B), and orientation bias appears clearly on rose plots in the direction of stripes, with an anisotropy index of 0.5 ± 0.1 for both ICAM-1 and VCAM-1 substrates. Although T cell can crawl and swim on adhesive and non-adhesive substrates, cells follow preferentially adhesive stripes, which suggest that crawling is more efficient than swimming.

**Figure 2:**
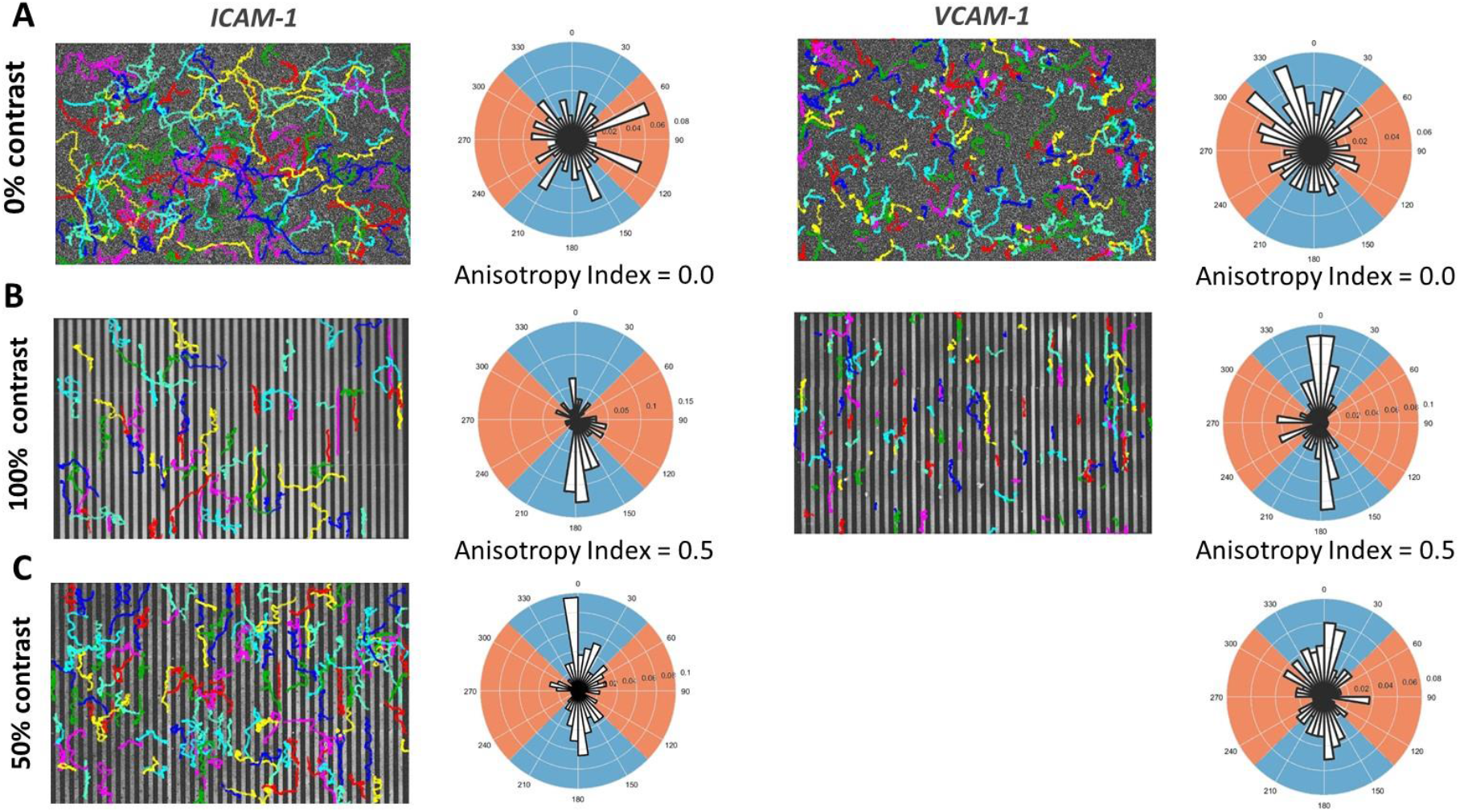
Substrates with modulated adhesion induce T cell haptotaxis. Cell tracking results on ICAM-1 (left panel) and VCAM-1 (right panel) substrates with modulated adhesion. **(A)** Negative control on substrates with 0% adhesion contrast where cells migrated in random directions. **(B)** Positive control substrates with 100% adhesion contrast where cells exhibited preferential haptotactic migration in the vertical quadrants (blue). **(C)** Substrates with 50% adhesion contrast where cells haptotaxed in the vertical quadrants (blue) with an intermediate anisotropy index between positive and negative controls. Each color represents the trajectory of migrating cell.

### Lymphocytes display integrin-mediated haptotaxis on globally adhesive substrates

Swimming is irrelevant for leukocyte recruitment from blood because firm adhesion on vessel walls is a prerequisite of transmigration to resist blood flow. We therefore investigated here cell orientation versus modulations of adhesion rate using stripes with different and non-null adhesion rates. **Figure 2**-C corresponds to patterns with alternated stripes of adhesion rates 0.6 and 0.3. To assess cell adhesion on each stripe type, we imaged the adhesion footprint of cells with the substrates by reflection interference contrast microscopy (RICM). **Figure 3** superimposes transmission images to localize cell body (grey), fluorescence microscopy images of the patterned stripes (red) and RICM images of the cell adhesion footprint (green). On homogeneously adherent substrates (**Figure 3**-A,D), cells display adherent lamellipods and non-adherent rear on ICAM-1, and conversely adherent rear and detached lamellipods on VCAM-1, as previously described(28). On substrates with adherent and non-adherent stripes (**Figure 3**-B,E), adhesion (green) signal was bright on integrin-ligand stripes (bright red) and extinct on anti-adhesive stripes (light red), which confirms that cells crawl on adherent stripes and swim over non-adherent stripes. For substrates with contrasted and non-null adhesion rates (**Figure 3**-C,F and see also suppl. Mat. Movie 3 and 4), RICM imaging confirms that cells adhere on both stripes types. These results confirm that cells are globally adhering on the substrates with stripes of modulated adhesion. The rose plots reveal nevertheless a clear preferential orientation in the direction of stripes with an anisotropy index of 0.3± 0.1 on both ICAM-1 and VCAM-1 (**Figure 2**-C). These data demonstrate unambiguously that lymphocytes are sensitive to modulations of substrate adhesion and develop integrin-mediated haptotaxis.

**Figure 3:**
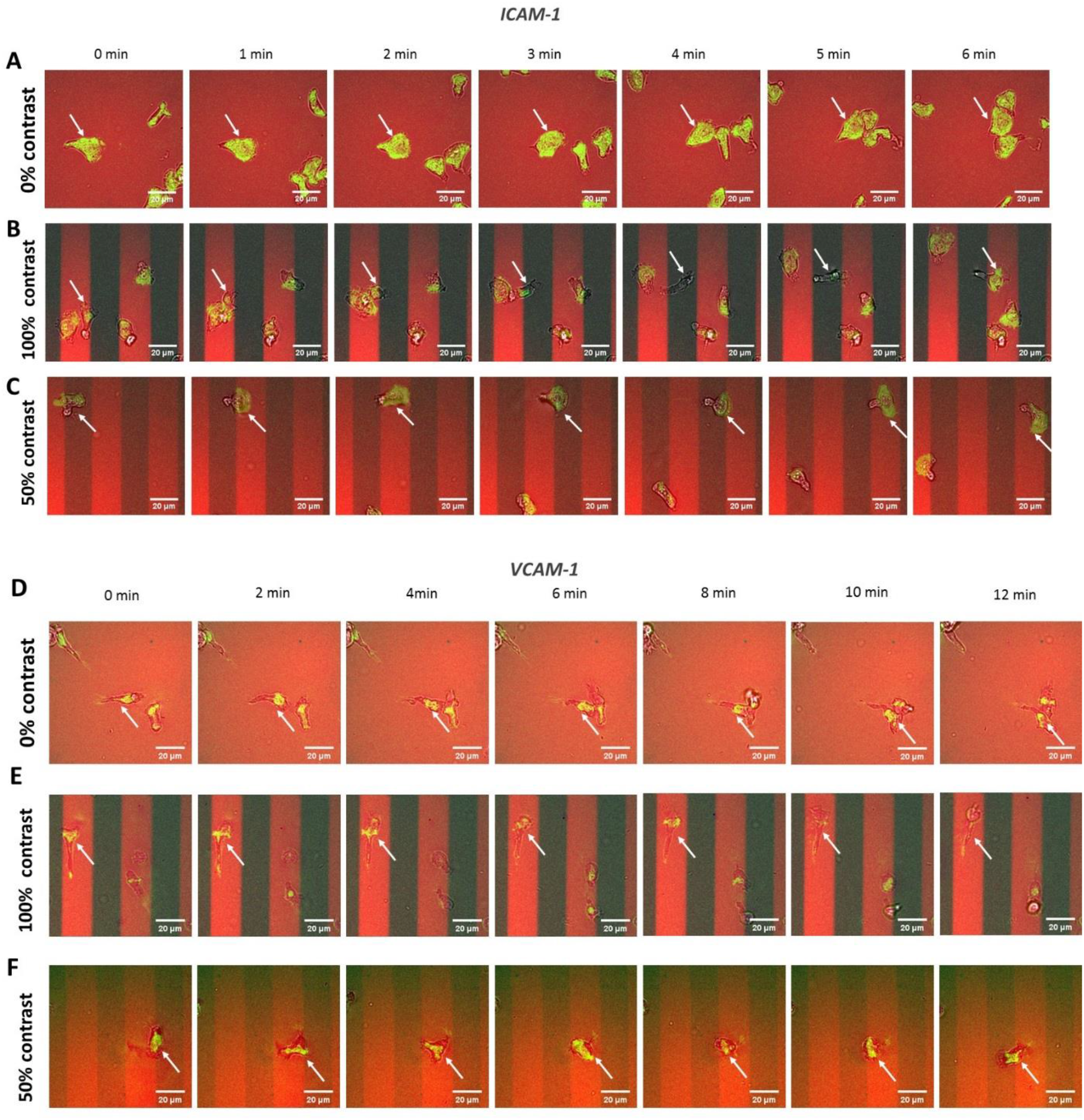
RICM shows different cell migration features on ICAM-1 and VCAM-1 substrates with modulated adhesion. **(A)** Negative ICAM-1 control substrate with 0% contrast shows cells migrating with fully spread lamellipods at the front. **(B)** Positive ICAM-1 control substrate with 100% contrast shows cells migrating either in or along adherent stripes or swimming across the non-adherent stripes. One can note the absence of RICM in between two adherent stripe, the large adhesion patch at cell front on adherent stripes and the general non-adherence of Cell uropod. **(C)** Substrate with intermediate contrast shows cell migrating across stripes with modulated adhesion. **(D)** Negative VCAM-1 control substrate with 0% contrast shows cells migrating with non-adherent lamellipods at cell front and attached middle cell body and uropod. **(E)** Positive VCAM-1 control substrate with 100% contrast shows cells migrating on adhesive stripe with attached cell rear and detached lamellipods. **(F)** Substrate with intermediate contrast shows cell migrating across stripes with modulated adhesion with attached cell rear and elevated lamellipod the entire time. In red: Fluorescent signal corresponding to integrin ligand concentration; In grey: bright-field image; In green: RICM adhesion footprint imaging.

### Efficiency of Integrin-mediated haptotaxis increases with adhesion contrast

To assess the range of molecular coatings allowing stimulation of integrin-mediated haptotaxis, we then varied the adhesion contrast of stripe-patterned substrates (**Figure 4**-A). The anisotropy index increased monotonously from 0 to 0.6 ± 0.2 when the adhesion contrast varied from 0 to 100%. The range in which substrates had modulated and non-null adhesion corresponded to adhesion contrast comprised within 0-65%. In these conditions, the anisotropy index reached a maximum of 0.46 ± 0.2. Interestingly, the crawling/swimming case, corresponding to adhesion contrast of 100%, appears as an extrapolation of the case of modulated adhesion. Furthermore, there is no apparent threshold for onset of haptotaxis versus adhesion contrast nor versus molecular surface concentration (Suppl. Mat. **Figure S 1**). These remarks suggest that haptotaxis may result from a similar mechanism with stripes of null and non-null adhesion.

**Figure 4:**
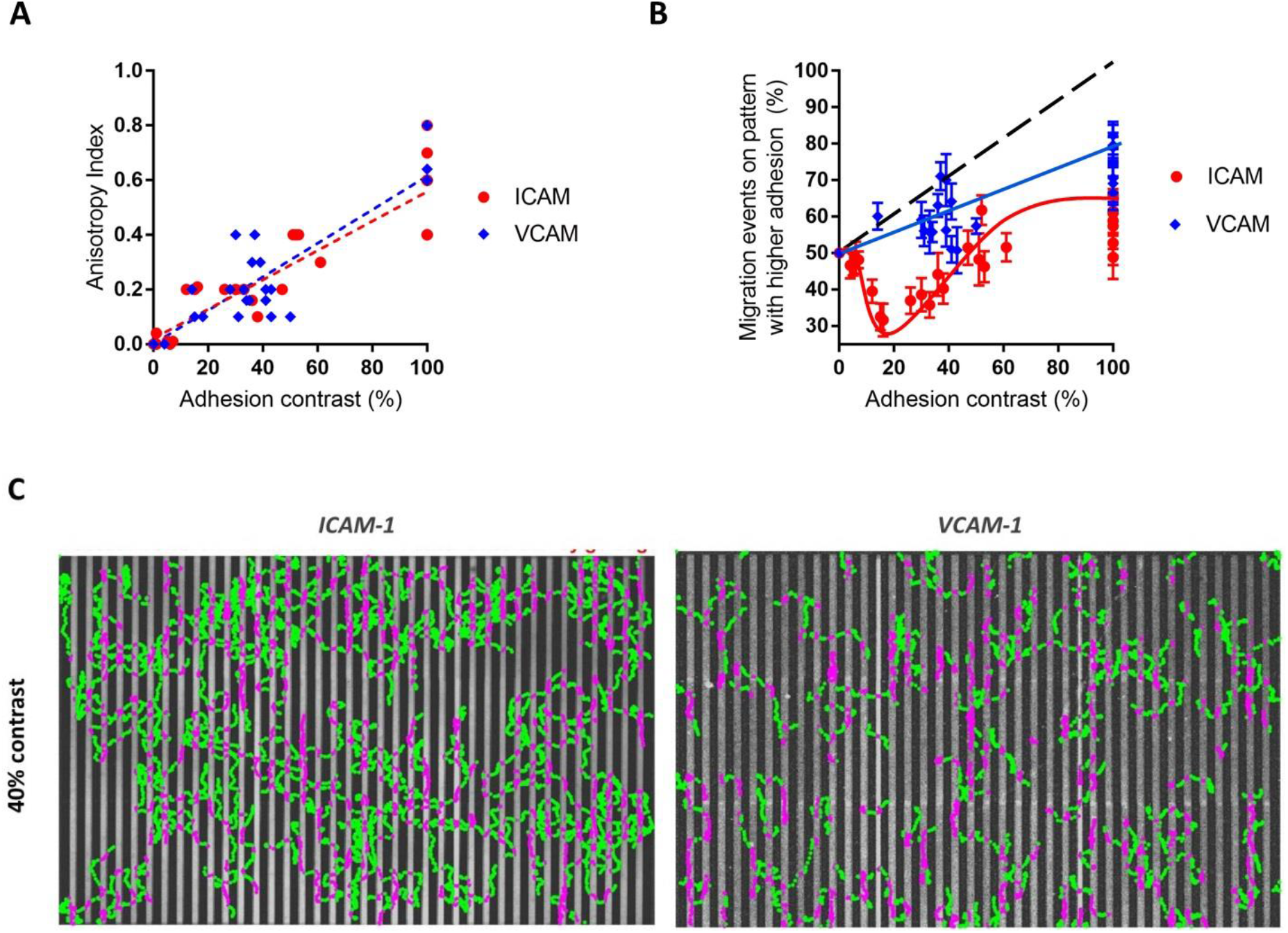
Integrin mediated haptotaxis displays different features on ICAM-1 and VCAM-1 substrates. **(A)** Anisotropy index versus adhesion contrast of ICAM-1 (red) and VCAM-1 (blue) substrates. A linear fit was used as a guide for the eye. Increasing adhesion contrast resulted in increasing anisotropy index with a slightly higher slope for VCAM-1 substrates. **(B)** Percentage of migration events on patterns with higher adhesion (MEHA) versus adhesion contrast of ICAM-1 (red) and VCAM-1 (blue). Solid lines are guides for the eye for both for experimental data whereas black dotted line is the trend line curve for mesenchymatous cells. **(C)** Characteristic cell trajectories on more adhesive (magenta) and less adhesive (green) ICAM-1 (left) and VCAM-1 (right) stripes with an intermediate adhesion contrast at 40%.

### VLA-4 integrins mediate a preference for high adhesion zones like mesenchymatous cells

To challenge further the role of simple adhesive steering, we then quantified the preference of cells between more or less adherent stripes by measuring the percentage of migration events on pattern with higher adhesion (MEHA) versus the adhesion contrast of the substrates (**Figure 4**-B). At adhesion contrast equal to 0, this percentage MEHA is 50 % for any cellular system because the difference between stripes vanishes. For lymphocytes on VCAM-1 substrates, the percentage MEHA increased monotonously from 50 % to 80 % versus adhesion contrast. A similar behavior is expected with mesenchymatous cell, as a tug of war mechanism favors a gradual increase of guiding versus the adhesion contrast experienced by cell edges. However, mesenchymatous must reach a percentage MEHA of 100 % for an adhesion contrast of 100%, because they are excluded from non-adherent zones. In contrast, lymphocytes reach only a maximum of 80%, which results from the capacity of amoeboid cells to swim over non-adherent surfaces. The common preference toward high adhesion zones for mesenchymatous cells in general and for lymphocytes on VCAM-1 substrates suggests the existence of a common mechanical bias by a passive tug of war mechanism.

### LFA-1 integrins mediate a unique preference for low adhesion zones

For lymphocytes on ICAM-1 substrates, the percentage MEHA versus adhesion contrast started also at 50% but then decreased down to a minimal value of 35% at lower adhesion contrast, whereas it systematically increased on VCAM-1 substrates. This surprising behavior reveals a marked preference of lymphocytes for less adherent zones on ICAM-1 coated substrates (**Figure 4**-B). Images of cell trajectories with a color-code to distinguish the fraction of cell paths on low and high adhesion stripes allowed us to evidence further a sharp difference between VCAM-1 and ICAM-1 cases (**Figure 4**-C). The counterintuitive preference for low adhesion zones observed on ICAM-1 substrates is clearly not compatible with a tug of war mechanism, and LFA-1 integrins mediate adhesive mechanotaxis with a distinct and original mechanism as compared to mesenchymatous cells. Besides, the fact that two different integrins mediate different haptotaxis phenotypes support that the mechanisms underlying integrin-mediated haptotaxis imply specific properties of integrins beyond their adhesion function, like their affinity regulation, their expression level, or their mechanotransduction competence.

### Mechanotransduction is not detected in integrin-mediated haptotaxis

To shed light on the implication of mechanotransduction by integrins during haptotaxis, we monitored intracellular calcium activity. **Figure 5** shows that calcium activity for cells crossing stripes of various adhesion remained negligible for both ICAM-1 and VCAM-1 coated substrates, whereas control with ionomycin showed an instant intense signal. Since calcium signaling is shared by many intracellular signaling pathways, these data support that mechanotransduction is not involved in integrin-mediated haptotaxis.

**Figure 5:**
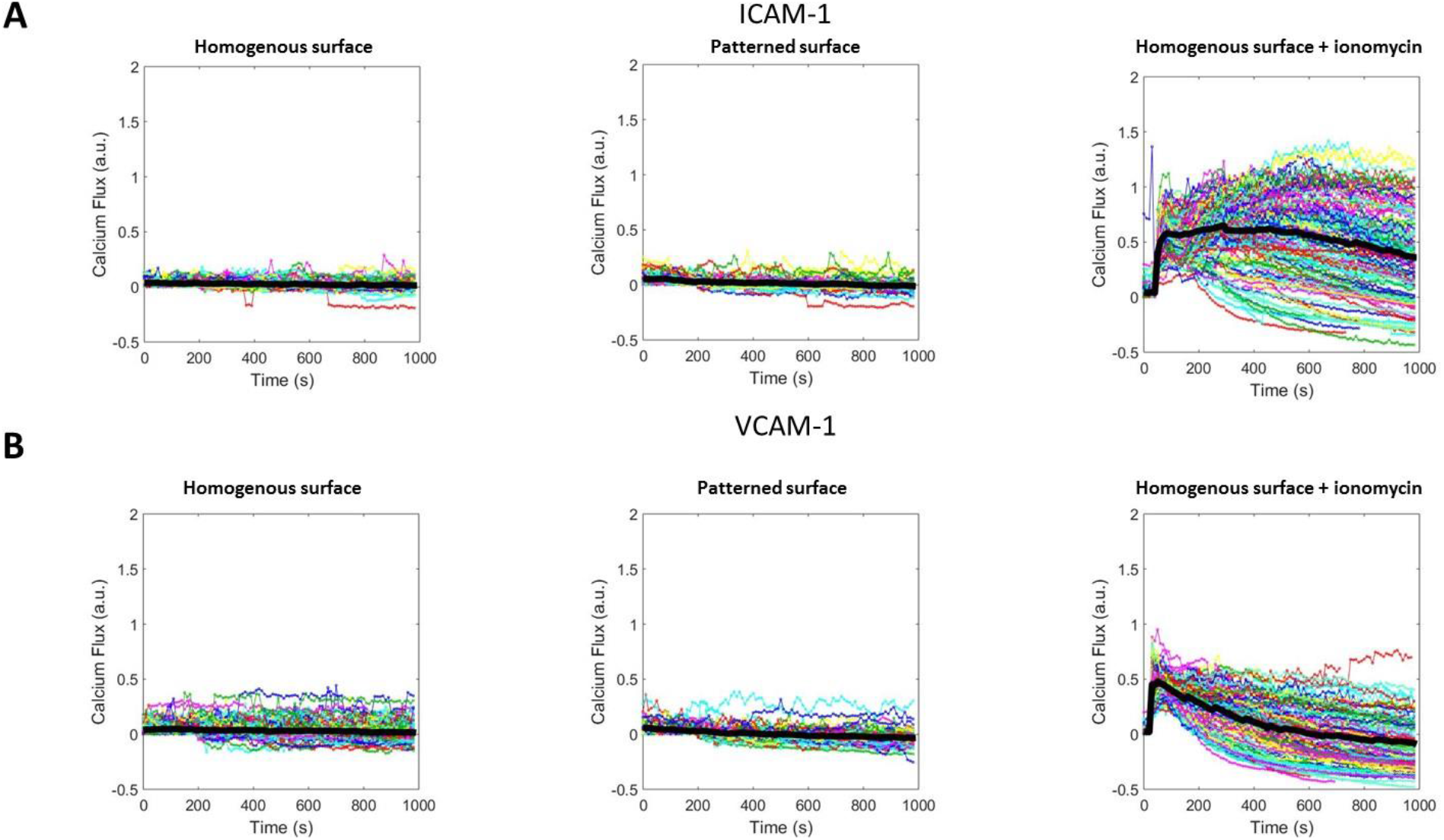
Intercellular Calcium flux detection reveals no mechanotransduction in integrin-mediated haptotaxis. The Calcium intensity was plotted as a function of time. Each colored line represents one migrating cell and the average intensity fluctuation was calculated and plotted with the thick black line. Calcium flux versus time on cells migrating on ICAM-1 **(A)** and VCAM-1 **(B)** substrates. On both homogenous surface (left) and surfaces patterned with stripes (middle), no intercellular Calcium flux was detected. The control experiment with ionomycin (right) revealed an instant strong signal.

## DISCUSSION

S.B. Carter(13) defined adhesive haptotaxis in 1965 with cancer cells migrating on cellulose acetate substrates and gradient of palladium. Cells moved robustly in the direction of increasing adhesion and Carter proposed that directed movement was controlled by the relative strength of their peripheral adhesion, hence the name ‘haptotaxis’ (Greek: haptein, to fasten; taxis, arrangement). Carter foresaw that movement towards surfaces offering greater adhesion may be a general phenomenon applicable to all metazoan cells, which are dependent on contact with a surface for their motility. Indeed, adhesive haptotaxis was later acknowledged for fibroblasts, neurons, stem cells(16–18, 29, 30). Repeatedly, adhesive steering was attributed to a tug of war between rival parts of the leading edge, those parts that have the strongest grip on the substrate being expected to win. In contrast, the migration of amoeboid cells, and in particular leukocytes, can take place without adhesive contact with a matrix (24) and even without a matrix (27). Directed motion controlled by adhesion is therefore not conceptually straightforward for amoeboid cells, and indeed, adhesive haptotaxis of leukocytes has, to our knowledge, never been evidenced. Our results here prove that lymphocytes are indeed sensitive to modulations of integrin-ligands concentration anchored on a matrix. Furthermore, the guiding effect observed in vitro on globally adhesive substrates may have a direct physiological relevance for the leukocytes crawling in blood vessels, because guidance was observed for ligand concentration modulated around 1000 molecules/μm^2^, which corresponds to expression of endothelial cells(31). Our results therefore validate the idea that overexpression of VCAM-1 and ICAM-1 at transmigration portals can engage crawling cells toward transmigration(3).

Our work shows also that adhesive haptotaxis differs strongly between lymphocytes, mesenchymatous cells or neuronal cells. Mesenchymatous cells like fibroblasts are totally excluded from non-adhesive zones and forced to follow by default the direction of adhesive stripes(30). Similarly, growth cones of neuronal cells avoid crossing from a more adhesive to a less adhesive substrate(17). In contrast, leukocytes can travel across adhesive and non-adhesive stripes with either ICAM-1 or VCAM-1. This capability implies that a tug of war effect in the integrin-ligand zone is not always efficient and that leukocytes may have had to develop alternative mechanisms to perform adhesive haptotaxis. On VCAM-1 substrates, lymphocytes prefer high adhesion zones, so that guidance across adhering and swimming zones appears as an extrapolation of guidance on substrates with adhesion modulations. Such properties can also be explained by a tug of war effect. In contrast, a tug of war mechanism is irrelevant for lymphocyte haptotaxis on ICAM-1 substrates, because cells show a predilection for lower adhesion zones. These results reveal a novel guidance mechanism mediated by integrins LFA-1. An hypothetical mechanism could be that LFA-1-mediated haptotaxis may function like chemokine mediated haptotaxis(32, 33), in which the mechanotransduction by integrins would work in place of the chemotransduction by chemokine receptors. However, calcium release experiments supported that no intracellular signaling was triggered by modulations of the adhesive stimulus (6). The sharp difference between haptotaxis mediated by integrins LFA-1 or VLA-4 may instead be linked to the strong differences in the dynamics of their spatial distribution and activation at the cell surface(28). Elucidation of these mechanisms would therefore benefit from precise characterization at single cell level of the strength of peripheral adhesion, and of the cartography of integrins localization and affinity state.

From a physiological point of view, we showed here that integrins can mediate lymphocyte haptotaxis on endothelial cells and that the differential preferences for low or high affinity zones between different integrins reveals novel sophisticated functions of integrins during the recruitment sequence toward transmigration portals. The unique phenotype induced by LFA-1 may for instance have a link with the particular importance (3) of its ligand ICAM-1 to favor transmigration.

## Materials and methods

### Cells and reagents

Whole blood from healthy adult donors was obtained from the Etablissement Francais du Sang. Peripheral Blood Mononuclear Cells (PBMC) were recovered from the interface of a Ficoll gradient (Eurobio, Evry, France). T cells were isolated with Pan T cell isolation Kit (Miltenyi Biotec, Bergisch Gladbach, Germany), then activated with antiCD3/antiCD28 Dynabeads (Gibco by Thermo Fischer Scientific, Waltham, MA) according to the manufacturer’s instructions. Cells were subsequently cultivated in Roswell Park Memorial Institute Medium (RPMI; Gibco by Thermo Fischer Scientific, Waltham, MA) 1640 supplemented with 25 mM GlutaMax (Gibco by Thermo Fischer Scientific, Waltham, MA), 10% fetal calf serum (FCS; Gibco by Thermo Fischer Scientific, Waltham, MA) at 37°C, 5% CO2 in the presence of IL-2 (50 ng/ml; Miltenyi Biotec, Bergisch Gladbach, Germany) and used 7 days after activation. At the time of use, the cells were >99% positive for pan-T lymphocyte marker CD3 and assessed for activation and proliferation with CD25, CD45RO, CD45RA and CD69 makers as judged by flow cytometry.

### Flow chamber preparation

Glass cover-slips (NEXTERION cover-slip, #1.5H Glass D263, SCHOTT Technical Glass Solutions, Jena, Germany) were first activated by air plasma (Harrick Plasma, Ithaca, NY, USA) for 5 min. Activated glass cover-slips were treated in gas phase with 3-Aminopropyltriethoxysilane (APTS; Sigma-Aldrich, St.Louis, MI) for 1 h and then heated for 15 min at 95°C on a heating plate. Sticky-Slides VI 0.4 (Ibidi GmbH, Martinsried, Germany) were then mounted on treated glass cover-slips. The prepared flow chambers were incubated sequentially at room temperature with an Alexa FluorTM 647 conjugate Protein A (Thermo Fischer Scientific, Waltham, MA) solution at 50 μg/mL for 1 h, a Bovine Serum Albumin (BSA; Sigma-Aldrich, St.Louis, MI) solution at 4% (w/v) for 15 min, and Fc-ICAM-1 or Fc-VCAM-1 (Intercellular Adhesion Molecule 1; Vascular Cell Adhesion Molecule 1, R&D system, Minneapolis, MN) solution at 10 μg/mL overnight at 4°C. The flow chambers were rinsed extensively with Phosphate Buffered Saline solution (PBS; Gibco by Thermo Fischer Scientific, Waltham, MA) after each incubation.

### Photo-patterning of adhesion molecules

We used an inverted microscope (TI Eclipse, Nikon, France) coupled to a UV laser source and a Digital Micromirror Device (Primo™, ALVEOLE, Paris, France)(26). Grey level Patterns were projected on ICAM-1 or VCAM-1-treated substrates in presence of a soluble photo-activator (PLPP™, ALVEOLE, Paris, France) to gradually degrade the proteins(34). Samples were then rinsed with PBS solution and passivated with 4% (w/v) BSA (Sigma-Aldrich, St.Louis, MI) for 15mins at room temperature.

### Cell adhesion and migration assay

Cells were seeded into the flow chamber at the concentration of approximately 1.5 million cells/mL and incubated for 10 min at 37°C. Cell migration was recorded at 37°C with a Zeiss Z1 automated microscope (Carl Zeiss, Oberkachen, Germany) equipped with a Snap HQ CCD camera (Photometrics, Tucson, AZ), pE-300 white LED microscope illuminator (CoolLED, Andover, UK) and piloted by μManager(35). Flow of prewarmed and CO2 equilibrated culture media through the flow chamber was controlled using an Ibidi pump system (Ibidi GmbH, Martinsreid, Germany). For cell adhesion assay, 20 bright-field images (Plan-Neofluar 10x/0.3 objective) were first collected every 10 s without flow. The flow chamber was then rinsed with culture media at 1 dyn.cm^−2^ to remove non-adherent cells. After rinsing, 100 bright-field images (Plan-Neofluar 10x/0.3 objective) were collected every 10 s without flow then under a shear stress at 4 dyn.cm^−2^ and finally under a shear stress at 8 dyn.cm^−2^. Fluorescent images for each pattern were collected at the end of the experiment with the same objective at recorded position. For cell migration assays, 20 bright-field images (Plan-Neofluar 10x/0.3 objective) were first collected every 10 s without flow. Then, the flow chamber was rinsed with culture media at 1 dyn.cm^−2^ to remove non-adherent cells. After rinsing, 100 bright-field images (Plan-Neofluar 10x/0.3 objective) were collected every 10 s. Fluorescent images for each pattern were collected at the end of the experiment with the same objective and at the same position. Additionally, bright-field and RICM images (Neofluar 63x/1.25 antiflex) were collected every 5 s for each pattern to reveal cell adhesion fingerprint, and fluorescent images were collected at the end of the experiment to localize the protein patterns.

### Fluorescent quantification of adhesion molecules

PE-labeled Anti-Human CD54 (ICAM-1) and Anti-Human CD106 (VCAM-1) antibodies (eBioScience by Thermo Fischer Scientific, Waltham, MA, USA) were used for adhesion molecules quantification. First, we set up a bulk calibration curve by measuring the fluorescence intensity of 41 μm thick channels filled with antibody solutions at concentrations of 1.5, 3, 5 and 7 μg/mL. Channels were pre-treated with 1% Pluronic F127© (Sigma-Aldrich, St Louis, MO) to limit adsorption of antibodies on channel surface. In the end, channels were rinsed extensively with PBS. Residual fluorescent intensity due to adsorbed antibodies was measured and then subtracted from the previous measurements. Previous study(28) showed a linear relation between the fluorescent intensity and the bulk concentration. We assume that the signal is given by the total number of molecules in the thin channel, then the volume concentration can be converted to a surface concentration for a channel of 41 μm in height. The surface of the flow chamber coated with ICAM-1 or VCAM-1 was then stained with corresponding antibody at 10μg/mL and incubated overnight at 4°C. Images were taken the next day with the Zeiss Z1 microscope set-up. The fluorescent intensity was analyzed with ImageJ software (U. S. National Institutes of Health, Bethesda, USA) at 5 different positions. The average intensity was converted into surface density of the adhesion molecules according the calibration data.

### Cell tracking and data analysis

A home-made program developed with MATLAB software (The MathWorks, Natick, MA) was used to track migrating cells as previously described in. Briefly, the program 1) performs image quality enhancement using background division and intensity normalization; 2) binarizes the images using a given threshold to distinguish cells from the background; 3) detects and numbers cells in the first frame and tracks them in the following frames and 4) saves the coordinates and time points for each identified cell to calculate different migration parameters. Only cells migrating for at least 30 μm are considered. In this study, to quantify cell adhesion, the program counts the number of adherent cells at 0 dyn.cm^−2^ and 8 dyn.cm^−2^ to determine the adhesion rate of patterned substrates. To determine the cell directionality on patterned surface, the migration angle is defined as the one between the patterned stripes and the total cell trajectory. The program calculates the fraction of cells with a migration angle <45° and >45°. The anisotropy index is defined as the difference between these two fractions. To further characterize the cell phenotypes on patterned surface, the program uses a binarized image of the fluorescent pattern as a mask to identify the position of each cell on the pattern. It then displays cell trajectories and calculates the percentage of migration events on patterns on higher or lower adhesion.

### Fluorescent detection of Calcium Flux

For calcium imaging experiments, cells were first seeded in channels with RPMI medium and were incubated for 10 min at 37°C to allow adhesion, then they were rinsed with HBSS + 1% BSA and incubated with Oregon Green^®^ 488 BAPTA-1, AM (ThermoFisher, Waltham, USA) diluted in HBSS + 1% BSA (5 μM) for 15 min at 37°C in the dark. After rinsing with HBSS + 1% BSA, the medium was replaced by HBSS+ 10% SVF. Control experiment was achieved by injection ionomycin (ThermoFisher, Waltham, USA) at a concentration of 1 μg.mL^−1^. The Calcium intensity fluctuation for each migrating cell was calculated for the whole duration of experiment as such: 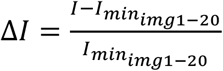. Then, the average fluctuation for all migrating cells was generated for the whole duration of experiment.

## Supplementary information

**Figure S 1: Anisotropy index versus surface density of integrin ligands**. For both ICAM-1 (in red) and VCAM-1 (in blue), the anisotropy index decreases with the surface density of integrin ligands.

**Figure S 2: Migration events on pattern with higher adhesion versus the adhesion contrast for immobile T cells**. On substrates with modulated adhesion, immobile cells adhere mainly on pattern with higher adhesion on VCAM-1 substrates (in blue) and on pattern with lower adhesion on ICAM-1 substrates (in red).

**Movie 1: T cell haptotactic migration on patterned ICAM-1 substrates**. Left: negative control substrate with 0% adhesion contrast; Middle: positive control substrate with 100% adhesion contrast; Right: substrate with 50% adhesion contrast. In red: substrate with higher adhesion; In grey: bright-field images of migrating cells.

**Movie 2: T cell haptotactic migration on patterned VCAM-1 substrates**. Left: negative control substrate with 0% adhesion contrast; Middle: positive control substrate with 100% adhesion contrast; Right: substrate with 50% adhesion contrast. In red: substrate with higher adhesion: In grey: bright-field images of migrating cells.

**Movie 3: RICM movies of T cells migrating on patterned ICAM-1 substrates**. Left: negative control substrate with 0% adhesion contrast; Middle: positive control substrate with 100% adhesion contrast; Right: substrate with 50% adhesion contrast. In green: cell adhesion patch shown by RICM; In red: substrate with higher adhesion; In grey: bright-field images showing cell body.

**Movie 4: RICM movies of T cells migrating on patterned VCAM-1 substrates**. Left: negative control substrate with 0% adhesion contrast; Middle: positive control substrate with 100% adhesion contrast; Right: substrate with 50% adhesion contrast. In green: cell adhesion patch shown by RICM; In red: substrate with higher adhesion; In grey: bright-field images showing cell body.

**Movie 5: Intracellular Calcium flux detection reveals no mechanotransduction on patterned ICAM-1 substrates**. Left: negative control substrate with 0% adhesion contrast; Middle: substrate with 50% adhesion contrast; Right: negative control substrate with ionomycine added.

**Movie 6: Intracellular Calcium flux detection reveals no mechanotransduction on patterned VCAM-1 substrates**. Left: negative control substrate with 0% adhesion contrast; Middle: substrate with 50% adhesion contrast; Right: negative control substrate with ionomycine added.

## Acknowledgements

The project leading to this publication has received funding from the ANR grant RECRUTE, the LABEX INFORM, the Région PACA, Institute CENTURI, Excellence Initiative of Aix-Marseille University –A*MIDEX, a French “Investissements d’Avenir” programme, and Alveole company. We are grateful to Laurence Borge for assistance with the use of the Cell Culture Platform facility (Luminy TPR2-INSERM).

## Bibliography

1. Massena, S., and M. Phillipso. 2012. Intravascular Leukocyte Chemotaxis: The Rules of Attraction. In: Lawrie C, editor. Hematology - Science and Practice. InTech.

2. Sumagin, R., H. Prizant, E. Lomakina, R.E. Waugh, and I.H. Sarelius. 2010. LFA-1 and Mac-1 Define Characteristically Different Intralumenal Crawling and Emigration Patterns for Monocytes and Neutrophils In Situ. J. Immunol. 185: 7057–7066.

3. Sumagin, R., and I.H. Sarelius. 2010. Intercellular Adhesion Molecule-1 Enrichment near Tricellular Endothelial Junctions Is Preferentially Associated with Leukocyte Transmigration and Signals for Reorganization of These Junctions To Accommodate Leukocyte Passage. J. Immunol. 184: 5242–5252.

4. Carman, C.V., and T.A. Springer. 2004. A transmigratory cup in leukocyte diapedesis both through individual vascular endothelial cells and between them. J. Cell Biol. 167: 377–88.

5. Valignat, M.P., O. Theodoly, A. Gucciardi, N. Hogg, and A.C. Lellouch. 2013. T lymphocytes orient against the direction of fluid flow during LFA-1-mediated migration. Biophys. J. 104: 322–331.

6. Valignat, M.-P., P. Nègre, S. Cadra, A.C. Lellouch, F. Gallet, S. Hénon, and O. Theodoly. 2014. Lymphocytes can self-steer passively with wind vane uropods. Nat. Commun. 5: 5213.

7. Dominguez, G.A., N.R. Anderson, and D.A. Hammer. 2015. The direction of migration of T-lymphocytes under flow depends upon which adhesion receptors are engaged. Integr Biol. 7: 345–355.

8. Gorina, R., R. Lyck, D. Vestweber, and B. Engelhardt. 2014. beta(2) Integrin-Mediated Crawling on Endothelial ICAM-1 and ICAM-2 Is a Prerequisite for Transcellular Neutrophil Diapedesis across the Inflamed Blood-Brain Barrier. J. Immunol. 192: 324–337.

9. Malawista, S.E., A. de Boisfleury Chevance, and L.A. Boxer. 2000. Random locomotion and chemotaxis of human blood polymorphonuclear leukocytes from a patient with Leukocyte Adhesion Deficiency-1: Normal displacement in close quarters via chimneying. Cytoskeleton. 46: 183–189.

10. Weber, M., R. Hauschild, J. Schwarz, C. Moussion, I. de Vries, D.F. Legler, S.A. Luther, T. Bollenbach, and M. Sixt. 2013. Interstitial Dendritic Cell Guidance by Haptotactic Chemokine Gradients. Science. 339: 328–332.

11. Canton, B. 2008. Refinement and standardization of synthetic biological parts and devices. Nat Biotechnol. 26: 787–793.

12. Schwarz, J., V. Bierbaum, K. Vaahtomeri, R. Hauschild, M. Brown, I. de Vries, A. Leithner, A. Reversat, J. Merrin, T. Tarrant, T. Bollenbach, and M. Sixt. 2017. Dendritic Cells Interpret Haptotactic Chemokine Gradients in a Manner Governed by Signal-to-Noise Ratio and Dependent on GRK6. Curr. Biol. 27: 1314–1325.

13. Carter, S. 1965. Principles of cell motility: the direction of cell movement and cancer invasion. Nature. 208: 1183.

14. Carter, S.B. 1967. Haptotaxis and the Mechanism of Cell Motility. Nature. 213: 256.

15. Carter, S.B. 1967. Haptotaxis and the Mechanism of Cell Motility. Nature. 213: 256–260.

16. Smith, J.T., J.K. Tomfohr, M.C. Wells, T.P. Beebe, T.B. Kepler, and W.M. Reichert. 2004. Measurement of Cell Migration on Surface-Bound Fibronectin Gradients. Langmuir. 20: 8279–8286.

17. O’Connor, T.P., J.S. Duerr, and D. Bentley. 1990. Pioneer growth cone steering decisions mediated by single filopodial contacts in situ. J. Neurosci. 10: 3935–3946.

18. Thibault, M.M., C.D. Hoemann, and M.D. Buschmann. 2007. Fibronectin, Vitronectin, and Collagen I Induce Chemotaxis and Haptotaxis of Human and Rabbit Mesenchymal Stem Cells in a Standardized Transmembrane Assay. Stem Cells Dev. 16: 489–502.

19. Brandley, B.K., and R.L. Schnaar. 1989. Tumor cell haptotaxis on covalently immobilized linear and exponential gradients of a cell adhesion peptide. Dev. Biol. 135: 74–86.

20. Klominek, J., K.-H. Robert, and K.-C. Sundqvist. Chemotaxis and Haptotaxis of Human Malignant Mesothelioma Cells: Effects of Fibronectin, Laminin, Type IV Collagen, and an Autocrine Motility Factor-like Substance1..

21. Aznavoorian, S., M.L. Stracke, H. Krutzsch, E. Schiffmann, and L.A. Liottã. Signal Transduction for Chemotaxis and Haptotaxis by Matrix Molecules in Tumor Cells..

22. Mccarthy, J.B., and L.T. Furcht. Laminin and Fibronectin Promote the Haptotactic Migration of B16 Mouse Melanoma Cells In Vitro..

23. Lämmermann, T., B.L. Bader, S.J. Monkley, T. Worbs, R. Wedlich-Söldner, K. Hirsch, M. Keller, R. Förster, D.R. Critchley, R. Fässler, and M. Sixt. 2008. Rapid leukocyte migration by integrin-independent flowing and squeezing. Nature. 453: 51–55.

24. Paluch, E.K., I.M. Aspalter, and M. Sixt. 2016. Focal Adhesion–Independent Cell Migration. Annu. Rev. Cell Dev. Biol. 32: 469–490.

25. Ricoult, S.G., T.E. Kennedy, and D. Juncker. 2015. Substrate-bound protein gradients to study haptotaxis. Front. Bioeng. Biotechnol. 3: 40.

26. Strale, P.-O., A. Azioune, G. Bugnicourt, Y. Lecomte, M. Chahid, and V. Studer. 2016. Multiprotein Printing by Light-Induced Molecular Adsorption. Adv. Mater. 28: 2024-+.

27. Aoun, L., P. Negre, A. Farutin, N. Garcia-Seyda, M.S. Rivzi, R. Galland, A. Michelot, X. Luo, M. Biarnes-Pelicot, C. Hivroz, S. Rafaï, J.-B. Sibarita, M.-P. Valignat, C. Misbah, and O. Theodoly. Mammalian Amoeboid Swimming is propelled by molecular and not protrusion-based paddling in Lymphocytes. BioRxiv MS ID BIORXIV2018509182..

28. Hornung, A., T. Sbarrato, N. Garcia-Seyda, L. Aoun, X. Luo, M. Biarnes-Pelicot, O. Theodoly, and M.-P. Valignat. A Bistable Mechanism Mediated by Integrins Controls Mechanotaxis of Leukocytes. BioRxiv ID BIORXIV2018509091ID..

29. Amarachintha, S.P., K.J. Ryan, M. Cayer, N.S. Boudreau, N.M. Johnson, and C.A. Heckman. 2015. Effect of Cdc42 domains on filopodia sensing, cell orientation, and haptotaxis. Cell. Signal. 27: 683–693.

30. Doyle, A.D., F.W. Wang, K. Matsumoto, and K.M. Yamada. 2009. One-dimensional topography underlies three-dimensional fibrillar cell migration. J. Cell Biol. 184: 481–490.

31. Dustin, M.L., and T.A. Springer. 1988. Lymphocyte function-associated antigen-1 (LFA-1) interaction with intercellular adhesion molecule-1 (ICAM-1) is one of at least three mechanisms for lymphocyte adhesion to cultured endothelial cells. J. Cell Biol. 107: 321 LP–331.

32. Woolf, E., I. Grigorova, A. Sagiv, V. Grabovsky, S.W. Feigelson, Z. Shulman, T. Hartmann, M. Sixt, J.G. Cyster, and R. Alon. 2007. Lymph node chemokines promote sustained T lymphocyte motility without triggering stable integrin adhesiveness in the absence of shear forces. Nat. Immunol. 8: 1076–1085.

33. Schwarz, J., V. Bierbaum, J. Merrin, T. Frank, R. Hauschild, T. Bollenbach, S. Tay, M. Sixt, and M. Mehling. 2016. A microfluidic device for measuring cell migration towards substrate-bound and soluble chemokine gradients OPEN. Nat. Publ. Group..

34. Pasturel, A., P.-O. Strale, and V. Studer. 2018. A generic widefield topographical and chemical photopatterning method for hydrogels..

35. Edelstein, A., N. Amodaj, K. Hoover, R. Vale, and N. Stuurman. 2010. Computer Control of Microscopes Using μManager. In: Current Protocols in Molecular Biology. Hoboken, NJ, USA: John Wiley & Sons, Inc. pp. 14.20.1–14.20.17.

